# Comparative genomics of a poinsettia-associated phytoplasma and functional validation of its SAP11-homologous effectors that induce plant branching

**DOI:** 10.1101/2025.10.17.683024

**Authors:** Shen-Chian Pei, Nian-Pu Li, Ting-Ting Li, Ya-Ching Yang, Ting-Hsuan Hung, Chih-Horng Kuo

**Author notes:** Correspondence: Ting-Hsuan Hung;, Chih-Horng Kuo. Email: Shen-Chian Pei, Nian-Pu Li, Ting-Ting Li, Ya-Ching Yang, Ting-Hsuan Hung, Chih-Horng Kuo.

## Abstract

Phytoplasmas are insect-transmitted plant pathogens that manipulate host development through secreted effector proteins. While they are notorious for causing agricultural losses, in the ornamental plant poinsettia (*Euphorbia pulcherrima*), phytoplasma infection is uniquely harnessed to induce the commercially desirable free-branching trait. However, the effectors responsible for this phenotype have remained unknown. To address this question, we sequenced and analyzed the genome of ‘*Candidatus* Phytoplasma pruni’ PR2021, a strain associated with the high-branching cultivar Princettia Pink. Comparative genomics confirmed its species assignment and revealed an unusual effector repertoire. PR2021 lacks most previously described effectors but encodes two distinct SAP11 homologs, a family of effectors known to induce plant branching. Genomic context analysis showed that one homolog is located within a potential mobile unit (PMU) and is nearly identical to the SAP11 of the distantly related ‘*Ca*. P. asteris’, while the other is located outside PMU regions and is divergent in sequence and predicted structure. Functional assays using *Agrobacterium*-mediated transient expression in *Nicotiana benthamiana* demonstrated that each homolog independently induced significant branching, whereas co-expression did not enhance the phenotype, indicating overlapping functions. These findings establish a direct connection between poinsettia branching and SAP11-homologous effectors, providing the first experimental evidence linking phytoplasma effector activity to this horticulturally important trait. This work expands understanding of phytoplasma effector diversity and mobility, while offering a functional framework for developing pathogen-free strategies to modulate ornamental plant architecture.

**IMPACT STATEMENT:** Phytoplasmas are uncultivated bacterial pathogens that reprogram host development through secreted effectors. While they are notorious for causing agricultural losses, phytoplasma infection is uniquely harnessed to induce the desirable free-branching trait in poinsettia, although the molecular basis has remained unresolved. Through analysis of the complete genome of ‘*Candidatus* Phytoplasma pruni’ PR2021, a strain associated with a high-branching cultivar, we identified two SAP11-homologous effectors with contrasting genomic and evolutionary contexts. One appears vertically inherited and divergent from previously characterized homologs, whereas the other is embedded in a potential mobile unit and likely acquired through horizontal transfer. Importantly, both homologs induce branching despite substantial sequence divergence. Taken together, this work advances understanding of phytoplasma genome evolution and effector diversity, while providing experimental evidence that links effector function to host developmental manipulation. Beyond its horticultural relevance, it illustrates how horizontal gene transfer and lineage-specific retention shape phytoplasma effector complements, offering a foundation for future efforts to dissect and re-engineer effector–host interactions.

**DATA SUMMARY:** All genome assemblies analyzed in this study were obtained from the National Center for Biotechnology Information (NCBI) Genome Database. The accession numbers are provided in Table S1.

## INTRODUCTION

Phytoplasmas are insect-transmitted plant pathogens that infect > 1,000 plant species (Lee et al. 2000; Hogenhout et al. 2008; Bertaccini et al. 2014; Namba 2019). These uncultivated bacteria are classified into more than 40 species in the genus ‘*Candidatus* Phytoplasma’ (‘*Ca*. P.’) (Wei & Zhao 2022; Bertaccini et al. 2022). Similar to other related lineages in the phylum *Mycoplasmatota*, phytoplasmas are characterized by a lack of cell wall, small cell size, and reductive genomes with low GC content (Gasparich et al. 2025). However, phytoplasmas are unique among *Mycoplasmatota* genera in their ability of manipulating the development programs of infected plants, causing symptoms such as stunting (shortening of internodes, reducing the size of leaves and flowers), witches’ broom (proliferation of stems and leaves), virescence (greening of flowers), and phyllody (abnormal development of floral parts into leaf-like tissues).

These abnormalities of host development are modulated through small secreted proteins produced by phytoplasmas known as effectors. Notable examples include: (1) SAP05, which induces witches’ broom and prolongs the host lifespan (Huang et al. 2021); (2) SAP06, which stunts vegetative growth and increases seed dormancy (Correa Marrero et al. 2024); (3) SAP11/SWP11, which alters the morphogenesis of leaves and roots, phosphate starvation response, and defense response (Sugio et al. 2011, 2014; Lu et al. 2014; Chang et al. 2018; Bai et al. 2009; N. Wang et al. 2018; Nan Wang et al. 2018; Chiu et al. 2025); (4) SAP54/PHYL1, which induces leaf-like flower development and promotes insect colonization on the plants (MacLean et al. 2011; Maejima et al. 2014; MacLean et al. 2014; Orlovskis et al. 2025; Orlovskis & Hogenhout 2016; Suzuki et al. 2023; Chiu et al. 2025); and (5) TENGU, which causes witches’-broom and stunting (Hoshi et al. 2009; Sugawara et al. 2013), as well as down-regulation of jasmonic acid and auxin pathways to result in plant sterility (Minato et al. 2014). It is worth noting that all of these effectors were initially characterized in ‘*Ca*. P. asteris’ and some were later studied in other ‘*Ca*. P.’ species. The presence/absence patterns regarding the homologous genes of these effectors are highly variable among different ‘*Ca*. P.’ species and strains (Huang et al. 2022).

The ability of manipulating plant development, coupled with their wide host range and geographic distribution, made phytoplasmas serious threats to agriculture (Lee et al. 2000; Bertaccini et al. 2014). Intriguingly, in one notable exception, their unique ability of modulating plant development is harnessed for the production of an ornamental plant. In this case, the bushy growth and dwarfism induced by phytoplasma infection are favorable traits of poinsettia (*Euphorbia pulcherrima*), resulting in many axillary shoots and “flowers” (modified leaves with bright red/pink colors called “bracts”) (Lee et al. 1997). This “free-branching” phenotype is important for large-scale production of vegetative cuttings and also increases the commercial value. Therefore, the poinsettia branch-inducing (PoiBI) phytoplasmas associated with such trait have been integrated into most commercial poinsettia cultivars. Reflecting this importance, the annual market value of potted poinsettias in the USA is approximately $157 million USD (National Agricultural Statistics Service 2021)

Recent studies of this poinsettia-phytoplasma system demonstrated that phytoplasma titer in stock plants was positively correlated with the branching phenotype of propagated cuttings, and higher phytoplasma loads were consistently detected in source leaves from the lower parts of the plant (Lee et al. 2021). In addition, expression of the poinsettia phenylalanine ammonia-lyase gene (*EpPAL*) varied among tissues and was negatively correlated with phytoplasma titer, suggesting that *EpPAL*-linked defense pathways may restrict pathogen growth (Lee et al. 2022). However, the exact phytoplasma effectors associated with the poinsettia branching phenotype remained unknown. Identifying these effectors is not only critical for explaining the poinsettia branching phenotype but also offers broader opportunities to link effector function with mechanisms of plant development. In the long term, such insights may enable the replacement of phytoplasmas, which can be highly variable, with breeding strategies or synthetic effector mimics to achieve desirable traits.

To address this question, we conducted metagenomic shotgun sequencing of the poinsettia cultivar Princettia Pink. The complete genome sequence of phytoplasma strain PR2021 associated with this cultivar was determined using a hybrid assembly method that combined Illumina and Oxford Nanopore Technologies (ONT) sequencing results. To provide early data access for the research community, the genome sequence of PR2021 has been made publicly available and published as a genome announcement (Pei et al. 2023). In this work, we present the branching phenotype comparison among poinsettia cultivars that led to the selection of Princettia Pink as the target of our sequencing effort, detailed analysis of the PR2021 genome and comparisons with other phytoplasmas, and experimental validation of two distinct SAP11 homologs from PR2021 that can induce branching phenotype in model plant *Nicotiana benthamiana*.

## METHODS

### Quantification of branching performance among poinsettia cultivars

To identify a PoiBI phytoplasma that is likely associated with a strong branching phenotype of its host, we compared the branching performance among nine poinsettia cultivars. These include Christmas Mouse, Luv U Pink, Noel, Pepride Red, Peterstar, Rose Star, Prima Red, Princettia Pink, and Princettia ROSA, all from the collection maintained at the Taoyuan District Agricultural Research and Extension Station, Shulin Substation (New Taipei, Taiwan). For each cultivar, 32 cuttings containing six to seven nodes were generated from maternal plants in May 2023. Wounding sites were treated with a phytohormone solution (1,500 mg/L indole-3-butyric acid and 500 mg/L 1-naphthaleneacetic acid) prior to planting in rockwool grow cubes to promote adventitious root formation. After new roots were formed, the plants were repotted into 5-inch pots, then maintained in a glass greenhouse at the Substation. After the plants grew to have more than 10 nodes, they were pinched to leave exactly 10 nodes. The branching performance was evaluated at three weeks after pinching. Newly formed branches with at least 0.5 cm in length were defined as effective branches.

### Phytoplasma genome analysis

The procedures of comparative genomics and phylogenetic analysis were largely based on those described in our previous studies (Chung et al. 2013; Huang et al. 2022; Cho et al. 2020). More detailed information is provided in the following sections. Unless stated otherwise, the methods were based on the cited references and the bioinformatic tools were used with the default settings.

For analysis of PR2021 genome, the annotated assembly was obtained from National Center for Biotechnology Information (NCBI) GenBank (accession GCA_029746895.1). The chromosome map was drawn using Circos v0.69-6 (Krzywinski et al. 2009). To identify its close relatives, we obtained all 272 phytoplasma genome assemblies available from NCBI as of March 1, 2025 (Table S1). Assemblies with N50 <10 kb were excluded to minimize the impact of highly fragmented genomes. Pairwise genome-wide average nucleotide identity (ANI) values were calculated using FastANI v1.33 (Jain et al. 2018). Strains sharing ≥90% ANI with PR2021 were retained for comparative analyses.

After identification of strains that are closely related to PR2021, we referred to the ‘*Ca*. P.’ phylogenetic trees based on 16S rRNA gene (Huang et al. 2023) and core genome (Huang et al. 2022) to select other representative species for comparative analysis at genus level. *Acholeplasma laidlawii* (GCA_000018785.1) was included as the outgroup. Homologous genes were identified using BLASTP v2.11.0 (Boratyn et al. 2013) with e-value cutoff set to 1e-15, followed by clustering of similarity result using OrthoMCL v1.3 (Li et al. 2003).

To infer core genome phylogeny, homologous genes that are conserved as single-copy in all genome analyzed were used. For phylogenetic inference, multiple sequence alignments were prepared using MUSCLE v3.8.31 (Edgar 2004) and visualized using JalView v2.11 (Waterhouse et al. 2009). Maximum likelihood phylogenies were inferred using PhyML v3.3 (Guindon & Gascuel 2003) and visualized using FigTree v1.4.4. PHYLIP v3.697 (Felsenstein 1989) was used for bootstrap analysis. For comparison of effector gene content, the homologous gene clustering result was examined manually based on annotation.

For comparative analysis of potential mobile units (PMUs), which are important for the evolution of phytoplasma effectors (Chung et al. 2013; Tokuda et al. 2023; Ku et al. 2013), methodology of PMU identification in PR2021 and selection of representatives for comparisons were based on that established previously (Huang et al. 2022). Briefly, based on the knowledge that intact PMUs can be up to 30 kb in size, we first identified genomic regions with at least four PMU core genes (i.e., *tra5*, *dnaB*, *dnaG*, *tmk*, *hflB*, *himA*, *ssb*, and *rpoD*) (Bai et al. 2006), each within ≤ 15 kb from the nearest neighbor. The PMU core genes located between two *tra5* homologs, which mark the PMU boundaries, were manually examined for their orientation to confirm that these genes belong to the same PMU. Visualization was performed using genoplotR v0.8.11 (Guy et al. 2010).

### Bioinformatic analysis of SAP11 homologs

The methods for multiple sequence alignment and phylogenetic inference were based on that described for core genome phylogeny in the previous section. For detailed examination of the protein sequences, signal peptide was predicted using SignalP v5.0 (Almagro Armenteros et al. 2019). Nuclear localization signal was predicted using two separate tools, with positive result from either one considered as valid, including WoLF PSORT (Horton et al. 2007) webtool (organism type set to “Plant”) and PredictNLS v1.0.20 (Cokol et al. 2000). Coiled-coil domain was predicted using COILS-WRAP (Lupas et al. 1991) with the settings “-m MTIDK -w 1”.

The method for protein structure prediction followed a recent study (Mirkin et al. 2025), which used AlphaFold2 (Jumper et al. 2021) via ColabFold (Mirdita et al. 2022). Prediction parameters included msa_mode: MMseqs2 (UniRef+Environmental), num_models: 5, num_recycles: 12, and stop_at_score: 100. For each protein, the model with the highest average predicted local distance difference test score among the five generated was selected for visualization using webtool Mol* 3D Viewer (Sehnal et al. 2021).

### Transient expression assays

For experimental investigation of putative effector genes, we performed transient expression assays in *Nicotiana benthamiana* (Shi et al. 2020; Hoshi et al. 2009). The candidates include two SAP11 homologs identified in the genome of ‘*Ca*. P. pruni’ PR2021 (locus tags: PR2021_2540 and PR2021_3970). The codon-optimized sequences, designed for *Nicotiana benthamiana* and without the secretory signal peptide, were synthesized by GenScript (Piscataway, NJ, USA). The synthesized sequences were cloned into a potato virus X (PVX) based expression vector pgR106 (Lu et al. 2003) using NEBuilder HiFi DNA Assembly (New England Biolabs). The resulting vectors were transformed into *Agrobacterium tumefaciens* strain GV3101 with pSOUP (Addgene 165419) via electroporation. The empty vector was included as the negative control.

For agroinfiltration in three-week-old *Nicotiana benthamiana* plants, the agrobacterial strains harboring the desired plasmids were cultured in lysogeny broth with rifampicin (50 µg/mL), gentamicin (10 µg/mL), and kanamycin (50 µg/mL) for 48 hours at 28 °C, harvested by centrifugation at 4,000 x g for 15 minutes, and resuspended in infiltration buffer (10 mM MgCl2, 10 mM MES, 100 μM acetosyringone, pH 5.7) with OD_600_ adjusted to 0.5. Three leaves of each plant were randomly selected to perform infiltration of 2 ml bacterial suspension. At three weeks after leaf infiltration, the number of side branches with at least three unfolded leaves for each plant was recorded. In each batch, five plants were used as biological replicates for each treatment. A total of three batches were performed during June-August, 2025 using plant growth chambers in Academia Sinica (Taipei, Taiwan). Methods for statistical tests and visualization followed that described for poinsettia branching performance.

### Statistical tests and visualization of branching counts

The statistical significance was evaluated using the Kruskal–Wallis test followed by Dunn’s post-hoc tests with Bonferroni correction for multiple comparisons, implemented with the packages stats v4.4.0 and dunn.test v1.3.6 in the R statistical environment v4.4.0 (R Core Team 2024). A non-parametric approach was chosen because pre-examination of branching count data with residual quantile–quantile plots and the Shapiro–Wilk test, implemented with the stats v4.4.0, raised concern about violations of normality assumptions in both experiments. Data visualization was performed using ggplot2 v3.5.1 (Wickham 2016), with all individual data points shown. The mean ± standard deviation was plotted for ease of interpretation, and letters assigned with multcompView v0.1-10 denote groups significantly different at p < 0.05.

## RESULTS AND DISCUSSION

### Strong branching of Princettia Pink motivated genomic characterization of its phytoplasma

In our branching performance evaluation, the Kruskal–Wallis test followed by Dunn’s post-hoc comparisons separated the nine cultivars into three overlapping significance groups (a-c) (Fig. 1). The four top-performing cultivars were assigned to the highest group (a), while Prima Red and Noel were assigned to the lowest group (c). The remaining three cultivars fell into intermediate overlapping groups (ab, abc, bc), reflecting that they were not significantly different from at least one higher and one lower cultivar. The highest-performing cultivar, Princettia Pink, developed 5.5 ± 2.0 effective branches at three weeks after pinching, which is notably higher than the ∼2.4–2.7 range observed for the three lowest performing cultivars.

**Fig. 1.**
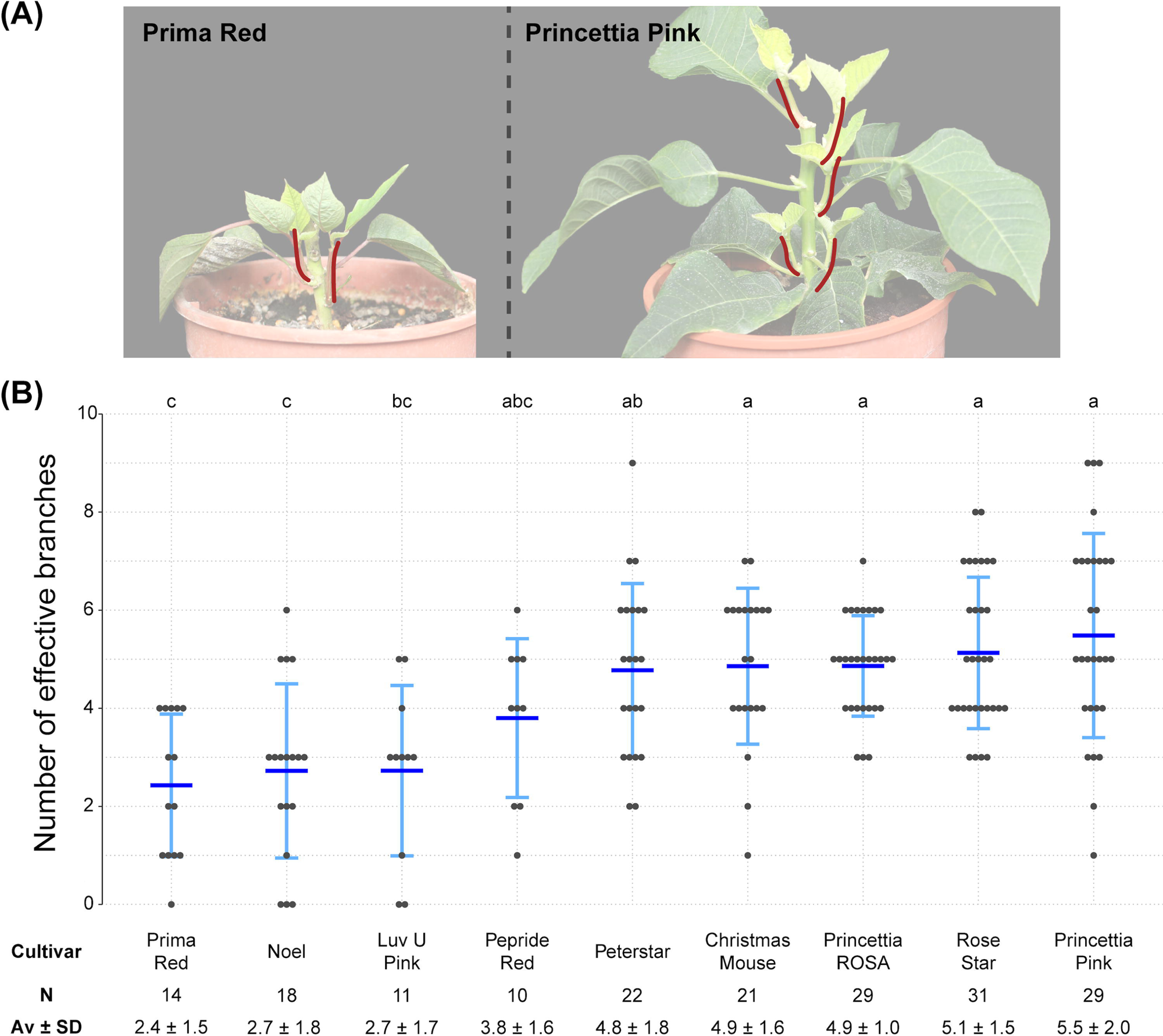
Branching performance among poinsettia cultivars. Poinsettia cuttings were pinched to exactly 10 nodes, and branching performance was evaluated three weeks later. Newly formed branches ≥0.5 cm in length were counted as effective branches. (A) Representative images of the low-branching cultivar Prima Red and the high-branching cultivar Princettia Pink. Effective branches are highlighted in red. (B) Quantification of branching performance across nine cultivars. Each point represents an individual plant. N indicates the number of cuttings scored per cultivar. Mean ± standard deviation are shown as horizontal bars in the plot and as numerical values below the x-axis. Statistical comparisons were performed using the Kruskal– Wallis test followed by Dunn’s post-hoc tests. Different letters indicate significant differences at p < 0.05 after multiple-testing correction.

Although only one batch was evaluated and seasonal effects were not tested, the performance difference was pronounced. Therefore, we selected Princettia Pink as the target for genomic characterization of its associated phytoplasma, designated strain PR2021. The vigorous growth of this cultivar also provided sufficient leaf midribs, the preferred tissue for metagenomic sequencing targeting phloem-residing phytoplasmas. As reported previously, we obtained the complete genome sequence of PoiBI phytoplasma strain PR2021 through a hybrid assembly approach (Pei et al. 2023).

### Comparative analysis of PR2021 and other phytoplasmas

For comparative analysis of PR2021 genome, we first identified its close relatives from publicly available assemblies (Table S1). Being uncultivated bacteria, phytoplasma genomes are typically reconstructed from metagenomic data. Due to the lack of reliable reference sets for assessing completeness and contamination, standard quality control tools such as CheckM (Parks et al. 2015) are not applicable. Overall assembly size is also unsuitable for evaluating completeness, since even complete circularized assemblies representing chromosomes within a single species can differ by nearly 50%, as the case of 576-853 kb range in well-studied ‘*Ca*. P. asteris’ (Cho et al. 2020). Therefore, we used assembly contiguity as a quality filter and excluded 49 highly fragmented assemblies with an N50 value below 10 kb. Considering the formal guideline of a 95% ANI threshold for delineating different ‘*Ca*. P.’ species (Bertaccini et al. 2022), we applied a more inclusive cutoff of 90% ANI to capture closely related but potentially distinct taxa, and identified eight strains (Fig. 2).

**Fig. 2.**
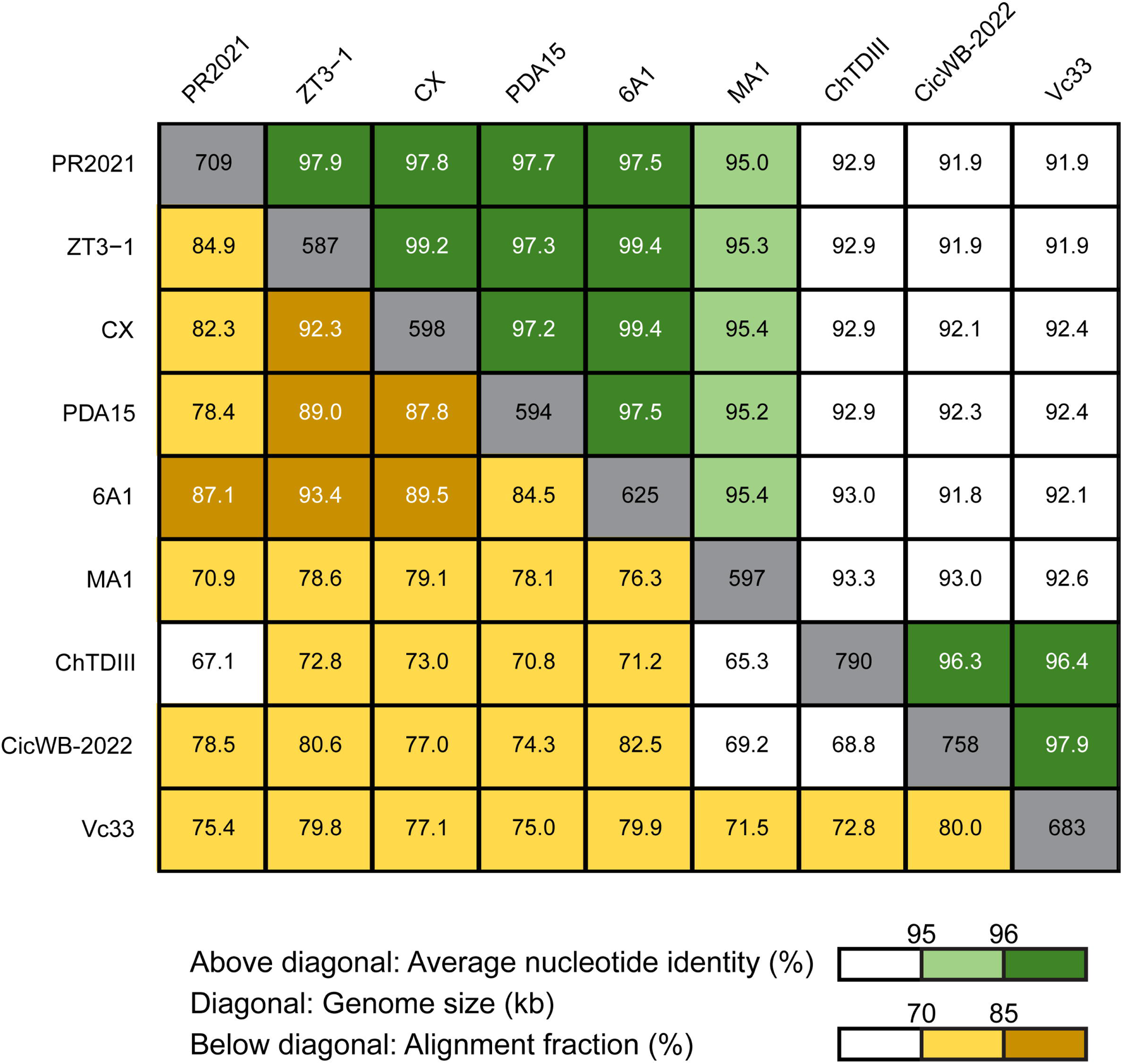
Pairwise genome similarity among ‘*Ca*. P. pruni’ PR2021 and closely related strains. The alignment fraction and average nucleotide identity (ANI) values for each pairwise comparison are provided below and above diagonal, respectively. Values in the diagonal cells indicate genome sizes.

Compared to these closely related strains, PR2021 shares 82.3% alignment fraction and 97.8% ANI with strain CX associated with Canada X-disease and described as ‘*Ca*. P. pruni’ (Lee et al. 2015; Davis et al. 2013), confirming its species assignment (Pei et al. 2023; Bertaccini et al. 2022). Meanwhile, although strain ChTDIII associated with chinaberry trees (*Melia azedarach* L.) in Argentina was initially identified as a member of ‘*Ca*. P. pruni’ (Fernández et al. 2020), this strain, together with two other South American strains, CicWB-2022 from Argentina (Fernández et al. 2024) and Vc33 from Chile (Zamorano & Fiore 2016), represent a distinct ‘*Ca*. P.’ species.

For genus-level analysis, we inferred a core genome phylogeny based on 163 conserved single-copy genes with 59,155 aligned amino acid sites (Fig. 3). Two PR2021-related strains, MA1 (197 contigs, N50 = 12 kb) and Vc33 (36 contigs, N50 = 36 kb), were excluded in the core genome analysis due to the lack of annotation in these assemblies. The phylogenetic placements further support the aforementioned species delineation, with strains with > 95% ANI forming distinct monophyletic clades. Comparative analysis of effector gene content revealed that PR2021 has two SAP11 homologs, but no other known phytoplasma effectors (Fig. 3). Given the complete circularized chromosomal contig of PR2021, these gene absences are unlikely to reflect false negative results caused by assembly artifacts.

**Fig. 3.**
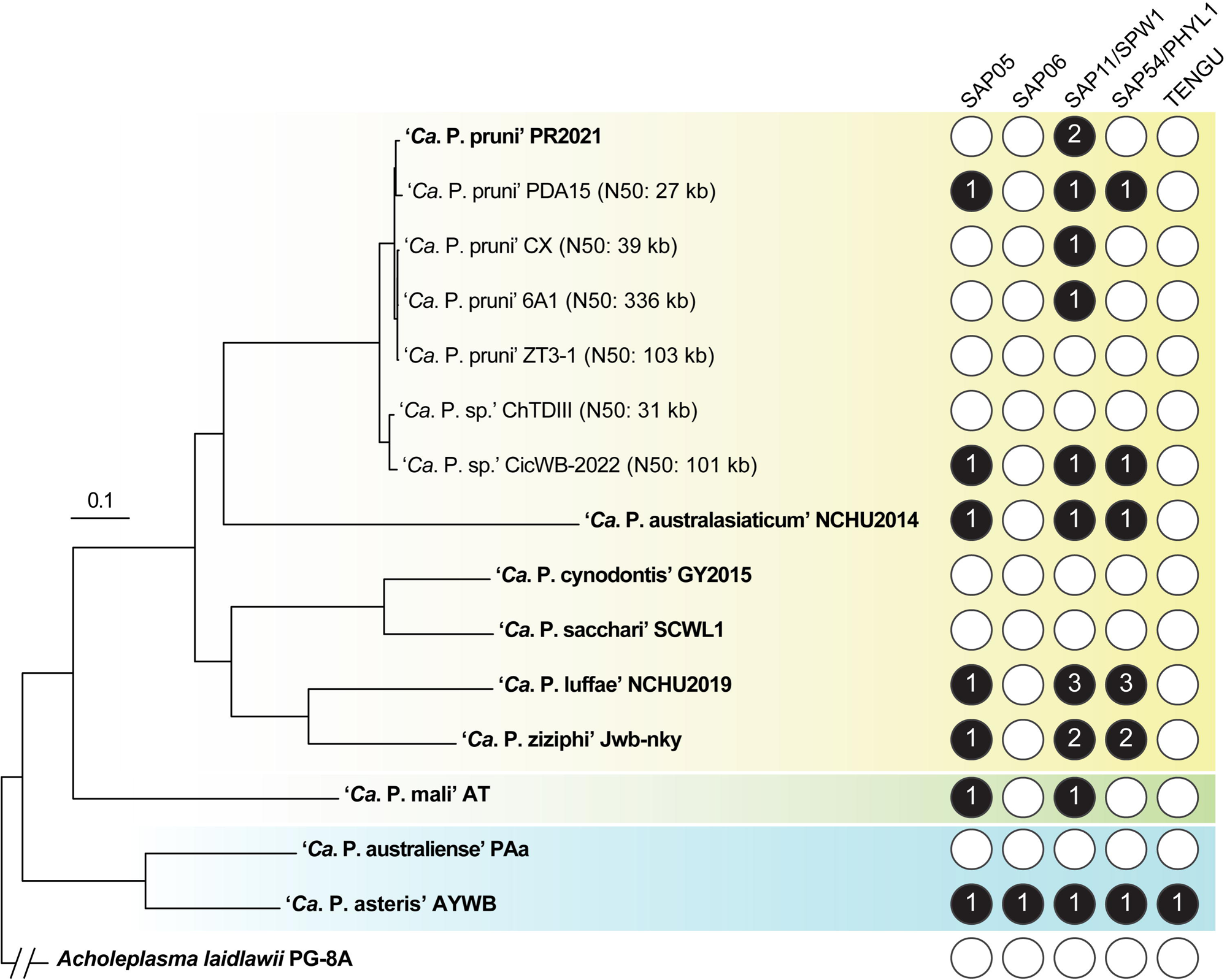
Core genome phylogeny and effector gene content. The maximum likelihood phylogeny was inferred using 163 conserved single-copy genes, the concatenated alignment contained 59,155 aligned amino acid sites. Based on 1,000 bootstrap resampling, the grouping of PR2021 and PDA15 received 61% support, all other internal nodes received > 98% support. Strains with complete genome assemblies are highlighted in bold, N50 values are provided in parentheses for strains with incomplete draft assemblies. The three major phylogenetic groups of phytoplasmas are indicated by colored background, *Acholeplasma laidlawii* was included as the outgroup. For the five characterized phytoplasma effectors, gene presence is indicated by filled circles, with copy number given inside; gene absence is indicated by empty circles.

This finding of effector gene content is noteworthy, because SAP11 homologs from diverse phytoplasmas have been reported to induce branching phenotypes. The effector SAP11 was first reported from ‘*Ca*. P. asteris’ strain AYWB and known to target plant nuclei (Bai et al. 2009). Transgenic expression of this gene in *Arabidopsis thaliana* increases stem numbers, reportedly through the destabilization of plant TEOSINTE-BRANCHED, CYCLOIDEA, PROLIFERATION FACTOR 1 and 2 (TCP) transcription factors (Sugio et al. 2011). Follow-up studies for transgenic expression of different phytoplasma SAP11 homologs in *Arabidopsis thaliana* (Chang et al. 2018; Nan Wang et al. 2018) and *Nicotiana benthamiana* (N. Wang et al. 2018) reported similar phenotypes and mechanisms. Based on these previous studies, the SAP11 homologs found in PR2021 are likely linked to the branching phenotype of its poinsettia host, therefore warrant further investigation.

### SAP11 homologs in PR2021 differ in genomic context and evolutionary origin

Intriguingly, the two SAP11 homologs in PR2021 have distinct genomic contexts. One homolog (locus tag: PR2021_2540) is 17.7 kb away from the nearest PMU gene and 69.0 kb away from the nearest intact PMU, therefore is not considered as PMU-associated (Fig. 4). By contrast, the other SAP11 homolog (PR2021_3970) is clearly located within a long PMU, designated as PR2021_2 (Figs. 4 and 5), suggesting distinct evolutionary histories for these two homologs.

**Fig. 4.**
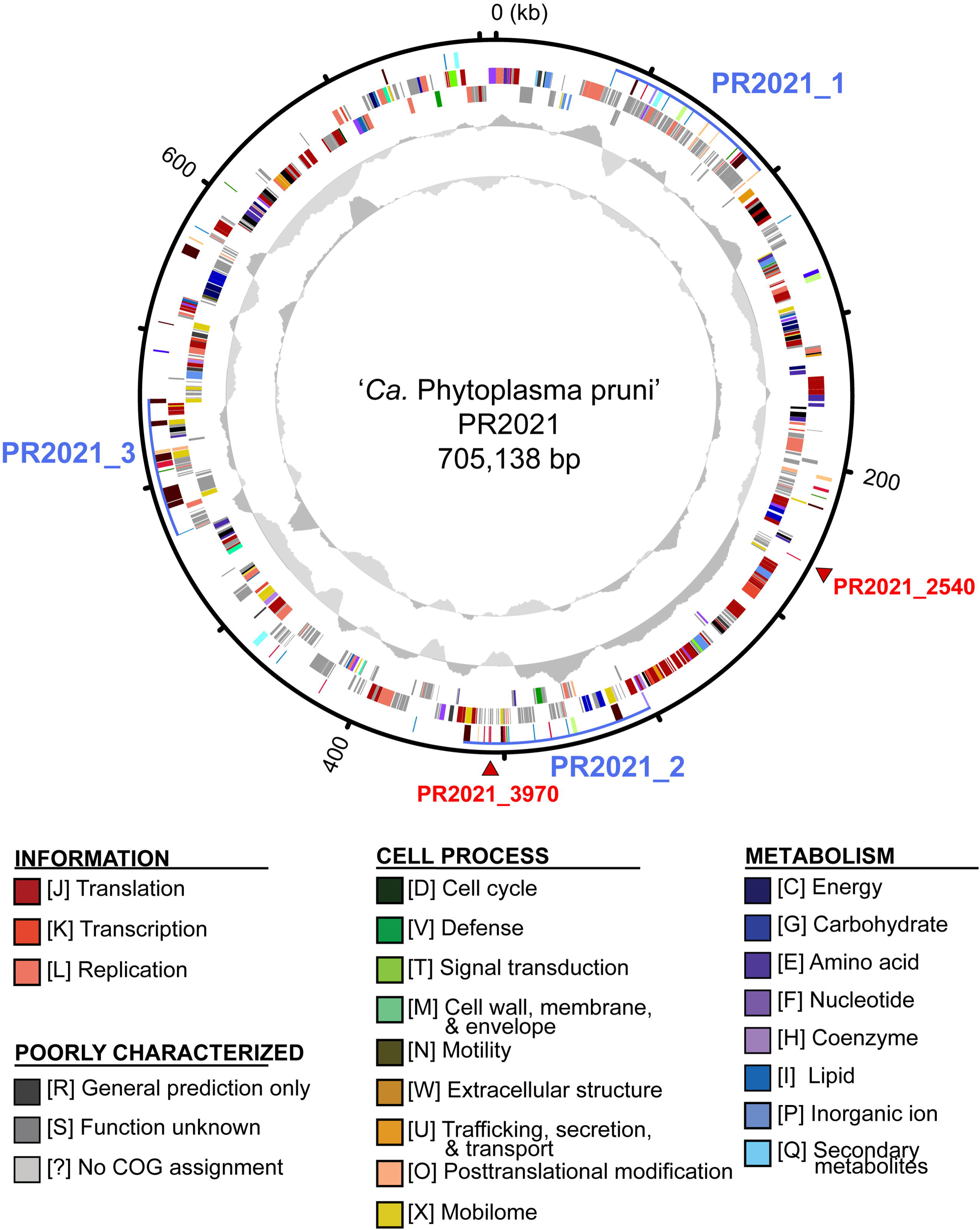
Chromosome map of ‘*Candidatus* Phytoplasma pruni’ PR2021. Concentric rings from outside in: (1) Scale marks (kb). (2) Genes associated with potential mobile units (PMUs) or annotated as encoding putative secreted proteins; color-coded according to the scheme illustrated in Fig. 5. The three gene clusters corresponding to intact potential mobile units (PMUs) are highlighted by blue lines. PMU identifiers and the two SAP11 homologs are indicated with blue and red labels outside of the scale mark ring, respectively. (3 and 4) Coding sequences on the forward and reverse strand, respectively. Color-coded by functional categories illustrated. (5) GC skew (positive: dark gray; negative: light gray). (6) GC content (above average: dark gray; below average; light gray).

**Fig. 5.**
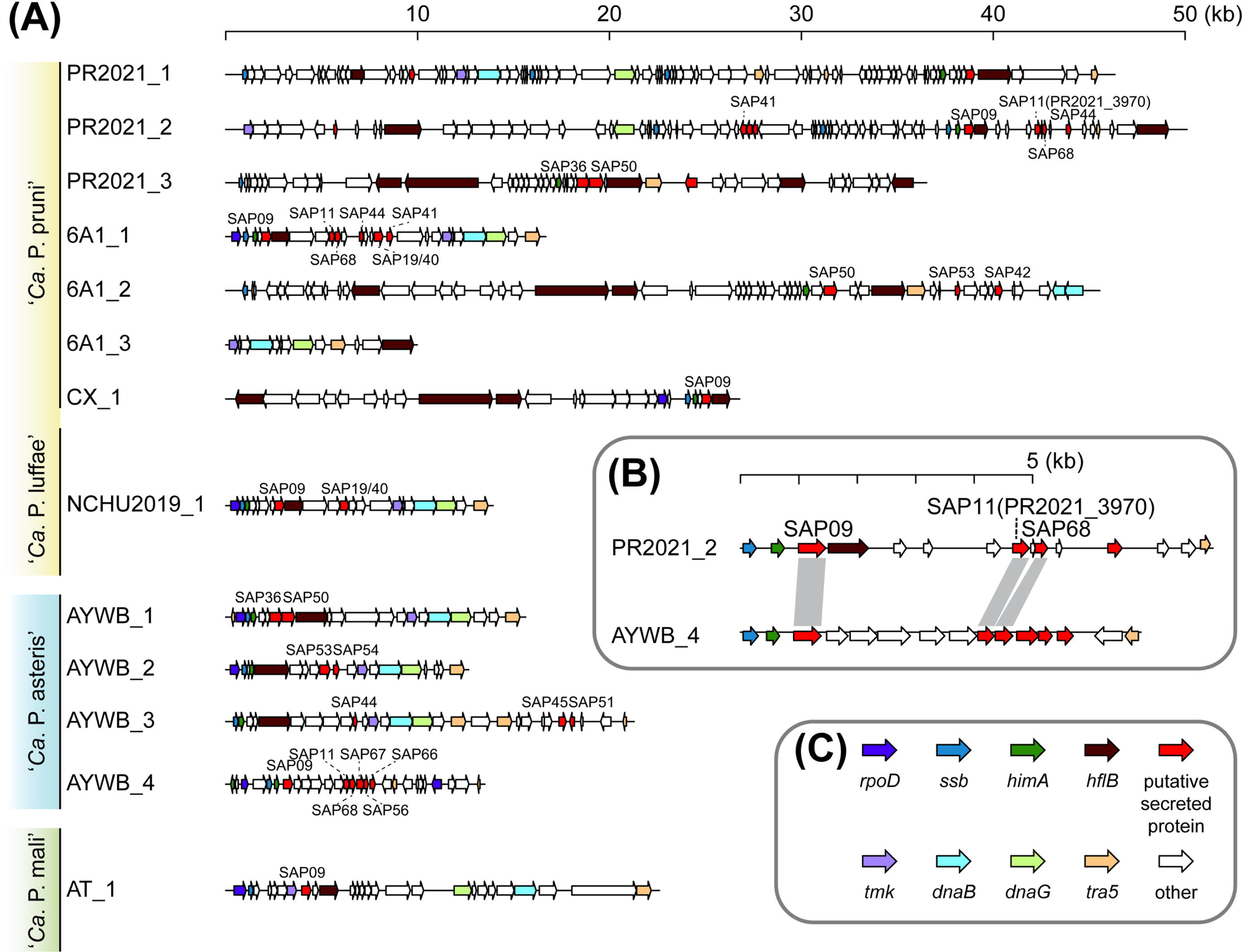
Gene organization of representative potential mobile units (PMUs). (A) Representative PMUs selected from diverse phytoplasmas. Defined PMUs are labeled with the phytoplasma strain name and a numerical identifier. Coding sequences in the defined regions were drawn to scale, with their orientations indicated by arrows. Homologs of previously described secreted AYWB proteins (SAP) are labeled. (B) An enlarged view comparing the regions containing SAP11 homologs between PR2021_2 and AYWB_4. (C) Color codes for PMU core genes and genes encoding putative secreted proteins.

Comparisons of PMUs among representative phytoplasmas revealed that ‘*Ca*. P. pruni’ harbors usually long PMUs. PR2021, which has a complete genome assembly, contains three intact PMUs ranging from 45 to 62 kb. Strain 6A1, which has a high-quality assembly with a N50 value of 336 kb, harbors three PMUs ranging from 12 to 56 kb. For strain CX, although only one 33 kb PMU was identified, PMU content could not be accurately assessed for this highly fragmented assembly with a N50 value of 39 kb. Together, compare to the size range of 6-28 kb observed in previously defined PMUs across diverse phytoplasmas (Huang et al. 2022) (Fig. 5), the newly characterized ‘*Ca*. P. pruni’ genomes revealed that phytoplasma PMU diversity is higher than previously recognized.

Our closer inspection of the PMU PR2021_2 revealed that PR2021_3970 is co-localized with several other SAP homologs, including SAP09 and SAP68 (Fig. 5B). A similar association among these genes was found in the PMU AYWB_4 from ‘*Ca*. P. asteris’, which contains the first experimentally characterized SAP11 (Bai et al. 2009; Sugio et al. 2011). Although PR2021_2 and AYWB_4 share little overall synteny conservation, possibly due to the long evolutionary distance between ‘*Ca*. P. pruni’ and ‘*Ca*. P. asteris’ (Fig. 3), the similarity of this SAP gene set is noteworthy and supports a role for PMUs in horizontal gene transfer across divergent phytoplasma lineages.

Sequence- and structure-based comparisons further highlight the contrasting evolutionary histories of the two PR2021 SAP11 homologs. The protein sequence encoded by PR2021_3970 shares 97.5% amino acid identity to ‘*Ca*. P. asteris’ SAP11 (locus tag: AYWB_370) (Fig. 6A) and clusters closely with it in phylogenetic analyses (Fig. 6B), despite the long divergence between the two species (Fig. 3). AlphaFold2 structural predictions also yielded nearly indistinguishable models (Fig. 6C). These observations are consistent with PR2021_3970 having been acquired recently through horizontal transfer, likely mediated by PMUs. By contrast, PR2021_2540 is highly divergent from AYWB_370, sharing only 39.0% identity and 47.9% similarity in their protein sequences (Fig. 6A). It shares 56.0-56.9% identity and 68.1% similarity to two other SAP11 homologs found in other ‘*Ca*. P. pruni’ strains, namely CPX_001780 from strain CX and QFY14_00375 from strain 6A1. These findings, together with the non-PMU chromosomal location of PR2021_2540, suggested that PR2021_2540 represents a vertically inherited homolog retained within ‘*Ca*. P. pruni’. This inference raises the question of whether these two evolutionarily distinct homologs are functionally equivalent.

**Fig. 6.**
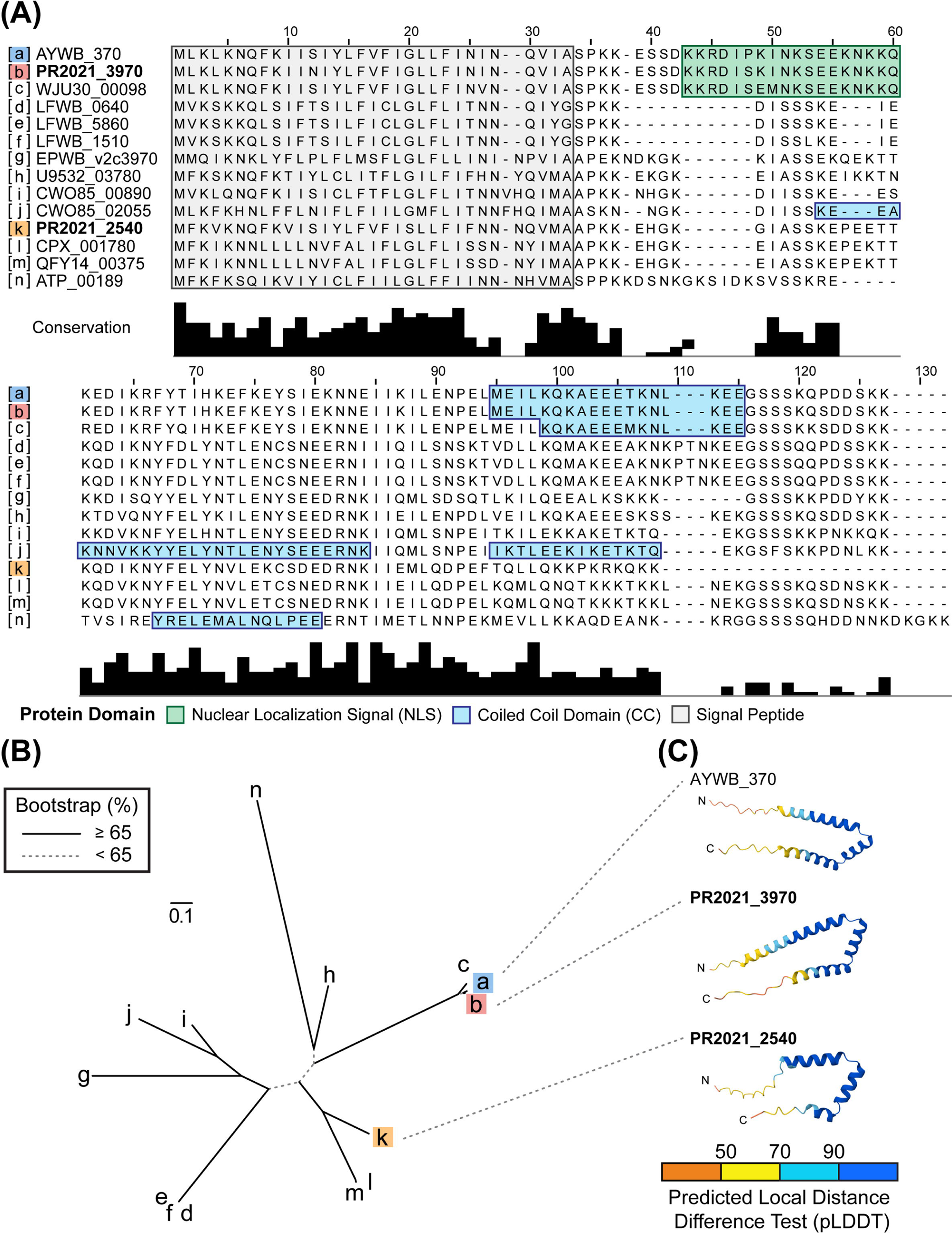
Bioinformatic analysis of SAP11 homologs. (A) Multiple sequence alignment. All homologs are identified by the locus tags from their respective genome annotation. The two homologs found in the PR2021 genome are highlighted in bold. Predicted domains are highlighted with colored backgrounds. (B) Maximum likelihood phylogeny. Bootstrap support levels were assessed based on 1,000 resampling. Homologs are labeled with lowercase alphabets as illustrated in panel A. (C) Protein structure prediction of the three key homologs, including the previously studied SAP11 from ‘*Ca*. P. asteris’ AYWB (locus tag: AYWB_370) and the two homologs identified in the PR2021 genome. Regions of the predicted structures are color-coded according to the confidence values based on predicted local distance difference test (pLDDT).

### Both SAP11 homologs are sufficient to induce branching in *Nicotiana benthamiana*

The findings that PR2021_3970 is nearly identical to the experimentally characterized AYWB_370, both in their amino acid sequences and predicted protein structures (Fig. 6), makes this SAP11 homolog an obvious candidate for explaining the branch-inducing property of PoiBI phytoplasmas. However, whether the second SAP11 homolog PR2021_2540 also plays a functional role is unclear. On one hand, it exhibits high levels of sequence divergence and bears lower similarity in the predicted protein structure. On the other hand, its predicted structure also contains three alpha helices as seen for PR2021_3970 and AYWB_370 (Fig. 6C). To empirically test if these two SAP11 homologs found in PR2021 are capable of inducing branching in host plants, we used a PVX-based transient expression system to express codon-optimized versions of these two genes, both individually and jointly, in *N. benthamiana*. At three weeks post agroinfiltration, control plants transformed with the empty vector were mostly unbranched, whereas expression of either homolog alone induced ∼4–5 side branches per plant, and co-expression averaged ∼6 branches (Fig. 7).

**Fig. 7.**
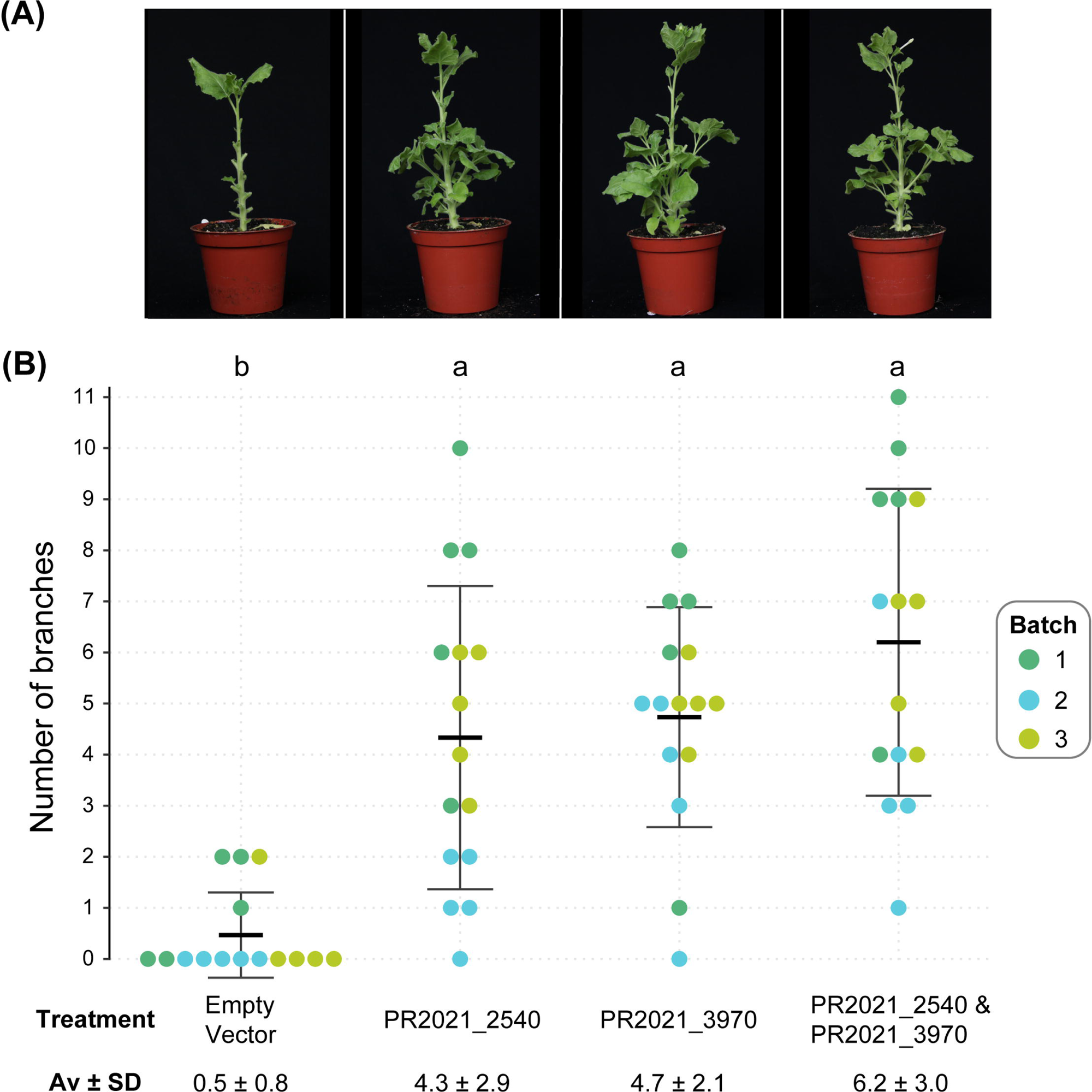
Experimental characterization of PR2021-encoded SAP11 homologs. SAP11 homologs were expressed in three-week-old *Nicotiana benthamiana* by agroinfiltration. Side branches with at least three unfolded leaves were counted three weeks post infiltration. Fully unfolded leaves were removed prior to imaging to facilitate branch counting. Each treatment included 15 plants in total, with five biological replicates per batch across three independent batches. (A) Photos of representative plants from each treatment. (B) Quantification of branch counts. Each point represents an individual plant. Mean ± standard deviation are shown as horizontal bars in the plot and as numerical values below the x-axis. Statistical comparisons were performed using the Kruskal–Wallis test followed by Dunn’s post-hoc tests. Different letters indicate significant differences at p < 0.05 after multiple-testing correction.

Although the expression levels of these genes were not directly quantified, consistent phenotypic differences were observed across independent batches. Rank-based comparisons showed that all three treatments differed significantly from the negative control, while not differing from one another. Thus, each SAP11 homolog is independently sufficient to induce branching. The activity of PR2021_3970 is consistent with its near identity to the AYWB SAP11 (Fig. 6) previously characterized in *Arabidopsis* (Sugio et al. 2011, 2014), whereas the comparable activity of the highly divergent, non-PMU PR2021_2540 is notable. The lack of significant additivity in co-expression suggests action on a shared host pathway, likely involving TCP destabilization, or saturation of the branching response, confirming functional redundancy. Together with the genomics and sequence analyses above, these results show that PR2021 harbors two distinct SAP11 homologs, allowing for robust manipulation of host development.

## CONCLUSION

In this study, we characterized the genome of the poinsettia-associated phytoplasma PR2021 and linked its effector repertoire to the free-branching trait of its host. Comparative genomics confirmed PR2021 as ‘*Ca*. P. pruni’ and revealed unusually long PMUs distinct from previously defined types. This genome encodes two SAP11 homologs, one likely retained through vertical inheritance and the other acquired through horizontal transfer, but no other known phytoplasma effectors. Functional assays demonstrated that each homolog is independently sufficient to induce branching in *N. benthamiana*. Notably, the branch-inducing property of PR2021_2540, despite its high sequence divergence from other SAP11 homologs, reveals functional conservation within this effector family that was not previously recognized. The absence of additive effects between the two homologs indicates functional redundancy, which may contribute to robust host manipulation. Beyond explaining an unusual plant– pathogen interaction with horticultural benefit, our results, together with prior work, highlight TCP transcription factors as promising targets for breeding or synthetic effector-based approaches to engineer branching traits without reliance on phytoplasma infection.

## AUTHOR STATEMENTS

### Author contributions

SCP performed poinsettia phenotyping, conducted all genomic and bioinformatic work, and generated Figures 1-6. NPL performed functional validation of effector candidates to prepare Figure 7. SCP and NPL both contributed to study design, data interpretation, and drafting the manuscript. TTL and YCY provided poinsettia cultivars, growth facility, assistance for maintaining plants, and horticultural knowledge. THH and CHK served as co-advisors for SCP and NPL, and provided expertise on poinsettia and phytoplasma biology. CHK conceived the study, acquired funding, administered the project, provided expertise on genomics, and wrote the manuscript. All authors read and approved the final manuscript.

### Conflict of interest

The authors declare no conflict of interest.

### Funding information

Funding was provided by Academia Sinica and the National Science and Technology Council of Taiwan (NSTC 113-2311-B-001-031) to CHK. The funders had no role in study design, data collection and interpretation, or the decision to submit the work for publication.

## Acknowledgements

We thank members of THH and CHK labs for discussion and assistance, Xiao-Hua Yan of the CHK lab helped with the preparation of Figure 1. Additional technical assistance was provided by Wei-Chun Gao (Taiwan Agricultural Research Institute, Ministry of Agriculture, Taiwan) and Hsin-Hung Yeh (Agricultural Biotechnology Research Center, Academia Sinica, Taiwan). The pgR106 vector was provided by David Baulcombe (University of Cambridge, UK).

## Usage of artificial intelligence tools

The authors used ChatGPT-5 to support brainstorming of manuscript organization, to suggest wording improvements, and to correct grammar. All substantive writing and the final text are the authors’ own work. No generative artificial intelligence tools were used in the preparation of figures or tables.

## Notes

### Competing Interest Statement

The authors have declared no competing interest.

